# The activity of chloroplast NADH dehydrogenase-like complex influences the photosynthetic activity of the moss *Physcomitrella patens*

**DOI:** 10.1101/2020.01.29.924597

**Authors:** Mattia Storti, Maria Paola Puggioni, Anna Segalla, Tomas Morosinotto, Alessandro Alboresi

## Abstract

Alternative electron pathways contribute to the regulation of photosynthetic light reactions to meet metabolic demands in a dynamic environment. Understanding the molecular mechanisms of their activity is seminal to decipher their role in response to environmental cues and in plant adaptation. The chloroplast NADH dehydrogenase-like (NDH) complex mediates cyclic electron transport pathway around photosystem I (PSI) in different organisms like cyanobacteria, algae and various plant species but has a discontinuous distribution in the green lineage. In order to assess how its activity and physiological role changed during plant evolution, we isolated *Physcomitrella patens* lines knocked out of the gene *NDHM* which encodes for a subunit fundamental for the stability and activity of the whole complex. *P. patens ndhm* KO mosses showed high PSI acceptor side limitation upon illumination leading to PSI photoinhibition. Flavodiiron proteins (FLV) have similar and particularly important role in preventing PSI overreduction when plants are exposed to light fluctuations. The *flva ndhm* double KO mosses alteration in photosynthetic parameters leaded to a defect in plant growth under fluctuating light as compared to WT and single KO mutants. Results evidenced that, while FLV sustain strong electron transport after an abrupt change in light intensity, NDH contribution to electron transport is small. NDH still participate in modulating PSI activity and it is seminal to prevent PSI photoinhibition especially when FLV are inactive. In plants the functional overlap between NDH- and FLV-dependent electron transport systems sustains PSI activity and to prevent its photoinhibition.

## INTRODUCTION

Photosynthetic reactions convert sunlight into chemical energy and sustain most lifeforms on earth. Light energy supports a linear electron flux (LEF) catalyzed by three major complexes imbedded in thylakoid membranes: (i) photosystem (PS) II, (ii) cytochrome b_6_f (Cyt b_6_f) and (iii) PSI. Overall electrons are transferred from water to NADP^+^ *via* plastoquinone (PQ), plastocyanin, ferredoxin and ferredoxin-NADP^+^ reductase. The electron transfer is coupled with proton (H^+^) accumulation in the lumen, originating from the photolysis of H_2_O and the reconversion of the reduced plastoquinone (PQH_2_) to PQ at the level of Cyt b_6_f. The LEF-generated H^+^ gradient is exploited by chloroplast ATPase for the synthesis of ≈1.3 ATP molecules per molecule of NADPH. Yet, the Calvin-Benson-Bassham (CBB) cycle is demanding ≈1.5 molecules of ATP per NADPH (Shikanai, 2016*a*). The cyclic electron flow (CEF) around PSI is one of the strategies adopted by photosynthetic organisms to divert electrons from reduced ferredoxin to reduce PQ, diminishing NADPH formation and increasing ATP biosynthesis (Arnon and Chain, 1975). Two major CEF pathways have been described so far although not conserved in all photosynthetic organisms (Peltier *et al.*, 2016; Yamori and Shikanai, 2016; Alboresi *et al.*, 2018). The first depends on PGR5/PGRL1 proteins (Munekage *et al.*, 2002; DalCorso *et al.*, 2008) and the second is mediated by chloroplast NADH dehydrogenase like complex (NDH-1, here called NDH), a multimeric complex which shares structural and evolutionary features with the mitochondrial respiratory complex I (Peltier *et al.*, 2016; Shikanai, 2016*b*). The NDH complex has a patchy distribution in the green lineage (Ruhlman *et al.*, 2015): it is present in cyanobacteria (Peltier *et al.*, 2016) and it has been identified in Gymnosperms, Angiosperms and also Streptophytae algae like *Klebsormidium flaccidum* (Hori *et al.*, 2014; Ruhlman *et al.*, 2015; de Vries *et al.*, 2016), but it is absent in some green algae like *Chlamydomonas reinhardtii* where a monomeric type II NADPH dehydrogenase (NDH-2; also called NDA2) was shown to play a role in CEF (Desplats *et al.*, 2009).

The NDH complex includes several subunits organized in various subcomplexes whose assembly is possible thanks to the activity of ancillary proteins. In angiosperms NDH forms a supercomplex with PSI through the binding with two specific antenna proteins, LHCA5 and LHCA6, as demonstrated by genetic and structural evidences (Peng *et al.*, 2008, 2009; Kouřil *et al.*, 2014; Yadav *et al.*, 2017). Further studies in *Arabidopsis* demonstrated that LHCA5 and LHCA6 replace respectively LHCA4 and LHCA2 in PSI for the binding of two PSI per NDH (Otani *et al.*, 2018). In the liverwort *Marchantia polymorpha*, which lacks the genes encoding for LHCA5 and LHCA6, PSI-NDH complex was not detected (Ueda *et al.*, 2012) suggesting that these subunits were acquired during the evolution of land plants possibly to allow a more efficient operation of NDH-dependent CEF pathway (Ueda *et al.*, 2012). In the moss *Physcomitrella patens*, where NDH complex is also present (Armbruster *et al.*, 2010; Kukuczka *et al.*, 2014) and active (Ito *et al.*, 2018; Kato *et al.*, 2018), the genome encodes for a *LHCA5* gene but not for *LHCA6* (Alboresi *et al.*, 2008) and NDH was suggested to associate with only one PSI complex (Kato *et al.*, 2018). Because of these peculiar structural properties and because of moss position in the tree of life, the functional investigation of NDH in *P. patens* can provide important insights on the function of alternative electron transports and how they were shaped by adaptation to terrestrial life. In fact, mechanisms for regulation of electron transport evolved and changed during plant evolution; for example, flavodiiron proteins (FLV) sustain an alternative pathway transferring electrons downstream PSI to O_2_ as a final acceptor with the formation of H_2_O. FLV activity has been reported in photosynthetic eukaryotes (Gerotto *et al.*, 2016; Shimakawa *et al.*, 2017; Chaux *et al.*, 2017; Jokel *et al.*, 2018) and in cyanobacteria (Allahverdiyeva *et al.*, 2013; Bersanini *et al.*, 2014), but they are not present in angiosperm genomes. In *P. patens* mutants lacking FLV their absence was suggested to be partially compensated by an increased CEF (Gerotto *et al.*, 2016) and showed complementarity with PGRL1/PGR5 cyclic electron transport (Storti *et al.*, 2019), but their functional interaction with NDH complex in plants has never been investigated so far.

We report the isolation of *ndhm* knocked-out (KO) mutants of *P. patens* and the resulting functional characterization of chloroplast NDH in this moss. Since *P. patens*, unlike angiosperms, has an active FLV-dependent electron transport pathway (Gerotto *et al.*, 2016), we investigated its functional overlap with NDH-dependent CEF pathway. Results evidenced that, while FLV sustain strong electron transport after an abrupt change in light intensity, NDH contribution to electron transport is small. Still NDH plays a role in modulating PSI activity and it is seminal to prevent PSI photoinhibition especially when also FLV are inactive.

## MATERIAL AND METHODS

### Plant Material and growth conditions

WT and mutant lines of *P. patens* (Gransden) were grown on PpNO_3_ medium at 24°C, 16 h light / 8 h dark photoperiod at control light intensity 50 μmol photons m^−2^s^−1^.

### Construct Design, Moss Transformation and Screening of Resistant

Regions from *NDHM* gene (Phypa_170083) were amplified by PCR from WT gDNA and cloned up- and down-stream the coding sequence of bleomycin resistance cassette (Fig. S1). This construct was used for PEG-mediated heat-shock protoplast transformation as previously described (Alboresi *et al.*, 2010). Two lines carrying one single insertion in *NDHM* target locus were named *ndhm* #1 and #2 and retained for further analysis. The *flva ndhm* double KO plants were selected on hygromycin-B by inserting *flva* mutation (Gerotto *et al.*, 2016) in *ndhm* #2. In all cases, stable mutant lines were isolated after two rounds of selection on antibiotic supplemented media. gDNA from resistant lines and control plants was extracted by a quick extraction protocol (Edwards *et al.*, 1991) and used to confirm the presence of the insertion at the expected target locus (Fig. S1; Table S1). 1 μg of total RNA purified with TRI Reagent (Sigma Aldrich) was used as template for cDNA synthesis with RevertAid Transcriptase (Thermo Scientific). *NDHM* gene expression was verified using specific primers and *ACT* gene was used as a reference (Fig. S1, Table S1) (Gerotto *et al.*, 2016). All experiments were performed using two independent lines for both *ndhm* and *flva ndhm* KO mutants. The average values obtained from the two lines are presented.

### SDS-PAGE, CN-PAGE and western blotting

Total Protein extracts were obtained by grinding fresh tissues directly in Laemmli buffer before loading SDS-PAGE. For immunoblotting analysis, after SDS-PAGE, proteins were transferred onto nitro-cellulose membrane (Pall Corporation) and detected with specific commercial (anti-PsaD, anti-γATPase and anti-Cyt f, Agrisera, catalog numbers AS09 461,AS08 312 and AS06119) or in-house made polyclonal antibodies (CP47, LHCA1, D2 and NDHM) (Bassi *et al.*, 1992). Thylakoid membranes for CN-PAGE were purified as in (Gerotto *et al.*, 2012) and resuspended in 25mM BisTris pH7, 20% glycerol buffer at final concentration of 1 μg chl/μL. Pigment-protein complexes were solubilized with 0.75% n-Dodecyl-α-Maltoside as described in (Järvi *et al.*, 2011) adding 0.2% deoxycholic acid prior loading. CN-PAGE gel were casted as described in (Järvi *et al.*, 2011) using a 3.5-11 % acrylamide gradient for isolation. Chl a/b and Chl/Car ratios were obtained by fitting the spectrum of 80% acetone pigment extracts with spectra of the individual pigments (Croce *et al.*, 2002).

### Fluorescence and P700 Measurement with Dual-PAM

Ten-day old plants grown on PpNO_3_ medium were probed for chlorophyll fluorescence and P700^+^ absorption with a Dual PAM-100 fluorometer (Walz). Plants were dark acclimated for 40 min before measurements. For light curves, light intensity increased every minute ranging from 0 to 230 μmol photons m^−2^ s^−1^. For induction kinetics, 50 or 540 μmol photons m^−2^ s^−1^ actinic red light was used, for fluctuating light kinetics a cycle of 3 min at 550 μmol photons m^−2^ s^−1^ and 9 min 25 μmol photons m^−2^ s^−1^ was repeated 5 times. PSII and PSI parameters were calculated as follow: Y(II), (Fm’-Fo)/Fm’; qL, (Fm’-F)/(Fm’-Fo’) x Fo’/F; NPQ, (Fm-Fm’)/Fm’; Y(I), 1-Y(ND)-Y(NA); Y(NA), (Pm-Pm’)/Pm; Y(ND), (1 - P700 red).

### Spectroscopic Analyses with Joliot-Type Spectrometer (JTS)

Spectroscopic analysis was performed *in vivo* on 10-day old intact tissues grown in control light using a JTS-10 spectrophotometer (Biologic). Electron transport rates (ETR) were evaluated measuring the Electrochromic Shift (ECS) spectral change on buffer infiltrated plants (Hepes 20mM pH 7.5, KCl 10mM) (Gerotto *et al.*, 2016). Relative amount of functional PSI was evaluated by xenon-induced single flash turnover in presence of 3-(3,4-dichlorophenyl)-1,1-dimethylurea (DCMU 20 μM) and hydroxylamine (HA, 4 mM) and used to normalize electron transport rate values. Because of the presence of double PSI turnovers using a xenon lamp (Bailleul *et al.*, 2010), ETR absolute values are underestimated by ≈40% (Gerotto *et al.*, 2016).

## RESULTS

### Isolation and biochemical characterization of Physcomitrella patens ndhm knock-out mutant

The functional role of *P. patens* chloroplast NDH complex was investigated by generating plants in which the gene encoding the NDHM subunit was knocked-out by homologous recombination. In *Arabidopsis thaliana* nuclear-encoded subunits of subcomplex A (e.g. NDHM, NDHN and NDHO) (Fig. 1) are known to be essential for the activity of the whole NDH complex (Rumeau *et al.*, 2005). Among them, NDHM was chosen as target for mutagenesis because in *P. patens* genome it is encoded by a single gene *locus* and it has already been detected in *P. patens* thylakoids by proteomic analysis (Kukuczka *et al.*, 2014). Two independent KO lines were confirmed by PCR to have a single insertion in the target *locus* and to have lost *NDHM* expression (Fig. S1). A clean deletion of the target gene was obtained thanks to the presence of direct LoxP recombination sites from the Cre/lox system that allowed the removal of the resistance cassette after transient expression of Cre recombinase in *ndhm* KO lines (Trouiller *et al.*, 2006). We analyzed the mutant lines by PCR amplification and sequencing of the entire recombinant locus (Fig. S1). The absence of NDHM protein was verified by western blotting using an antiserum raised against the full-length recombinant PpNDHM protein. Immunoblot analysis revealed that the accumulation of NDHH, a plastid-encoded subunit of subcomplex A, was also decreased by more than 90% in *ndhm* KO lines (Fig. 1B). This observation suggests that the absence of NDHM caused the destabilization of NDH subcomplex A as observed in *A. thaliana* (Peng *et al.*, 2008). The *ndhm* KO plants grown under optimum conditions did not show any major morphological alteration as compared to the WT (Fig. 1). Pigment composition also was unchanged in both *ndhm* KO lines (Table S2). Moreover, the *ndhm* KO mutants showed no significant difference in abundance as regards PSI and PSII core subunits (PsaD and D2) nor Cyt f and γ-ATPase (Fig. 1B).

**Fig. 1.**
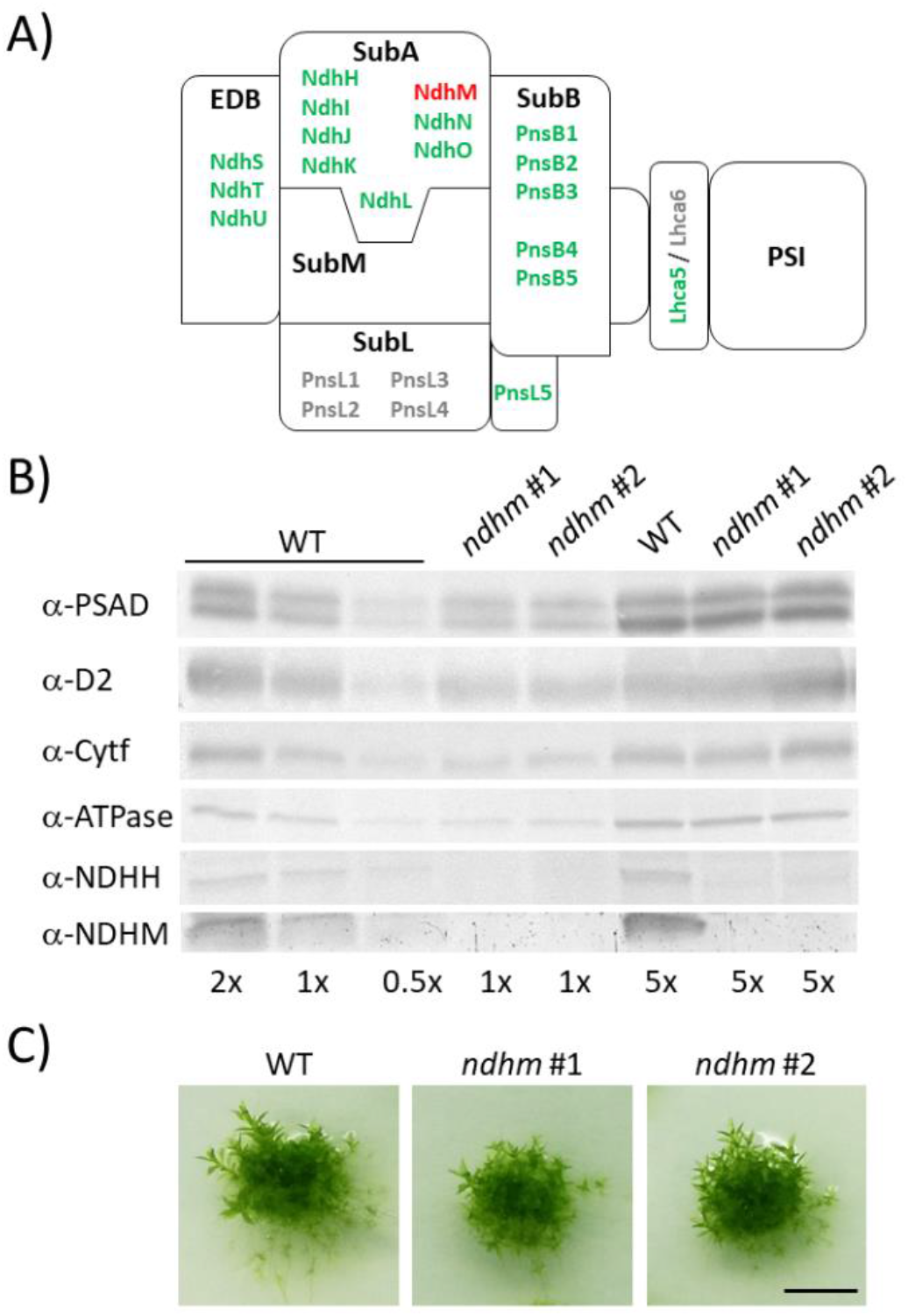
Phenotype of *ndhm* knockout (KO) mutants. A) Predicted structure of photosystem I (PSI)– NAD(P)H dehydrogenase (NDH) supercomplex of the model plant *Arabidopsis thaliana*. PSI is connected to NDH through the antenna linkers LCHA5/LHCA6. Different sub-complexes form NDH complex, namely EDB, SubA, SubB, SubL and SubM. All the sequences either putatively present or absent in *P. patens* genome are respectively labeled in green or grey. NDHM subunit is labeled in red. B) Immunoblot analysis for detection of PSAD, D2, Cytf, γ-ATPase, NDHH and NDHM in WT and *ndhm* KO #1 and #2 plants. 1X is equivalent to 2 μg of chlorophylls for the detection of D2 and NDHH, 3 μg of chlorophylls for the detection of PSAD, ATPase and NDHM and 4 μg of chlorophylls for the detection of Cytf. 5X, 2X and 0.5X is respectively five, two and half times 1X. C) Moss colonies of WT and *ndhm* KO #1 and #2 grown for 21 days on PPNO_3_ under long-day photoperiod 50 μmol photons m^−2^ s^−1^. Scale bars = 0,5 cm.

Protein complexes from the thylakoid membrane were separated using a clear native (CN) gel after a mild solubilization (Fig. 2). There was no detectable difference between WT and *ndhm* KO in the pattern of chlorophyll-containing green bands or in the protein bands after staining with Coomassie Brilliant Blue (Fig. 2A). CN-PAGE confirmed that under optimum growth conditions the photosynthetic apparatus of *ndhm* KO plants is very similar to the WT and in *P. patens* CN-PAGE profile no band was clearly identifiable as PSI-NDH supercomplex. This is different from *A. thaliana* and *Hordeum vulgare*, where a large supercomplex formed by one chloroplast NDH surrounded by two PSI has been characterized (Peng *et al.*, 2009; Kouřil *et al.*, 2014) and was detectable by native gel electrophoresis (Peng *et al.*, 2009; Järvi *et al.*, 2011). Proteins in CN-PAGE were separated after denaturation in a second dimension by SDS-PAGE and analyzed by immunoblotting using specific antibodies (Fig. 2B). Anti-NDHM antibody revealed the presence of NDH complex in two major spots, absent in the *ndhm* KO (Fig. 2B). The weaker spot corresponded to a native size of ≈700 KDa, as expected for NDH monomers and described in *M. polymorpha* (Ueda *et al.*, 2012). A more intense NDHM signal was instead detected at higher MW (≈1000 KDa in the native gel), comigrating with PSI and PSII supercomplexes (Fig. 2B). This spot corresponds to the approximate size of a PSI-NDH supercomplex with a single NDH and PSI as recently suggested (Otani *et al.*, 2018). An immunoblot analysis using anti-PSAD and anti-LHCA1 antibodies failed to reveal PSI subunits in the same region, suggesting that putative PSI-NDH represents a very small fraction of PSI particles in *P. patens*.

**Fig. 2.**
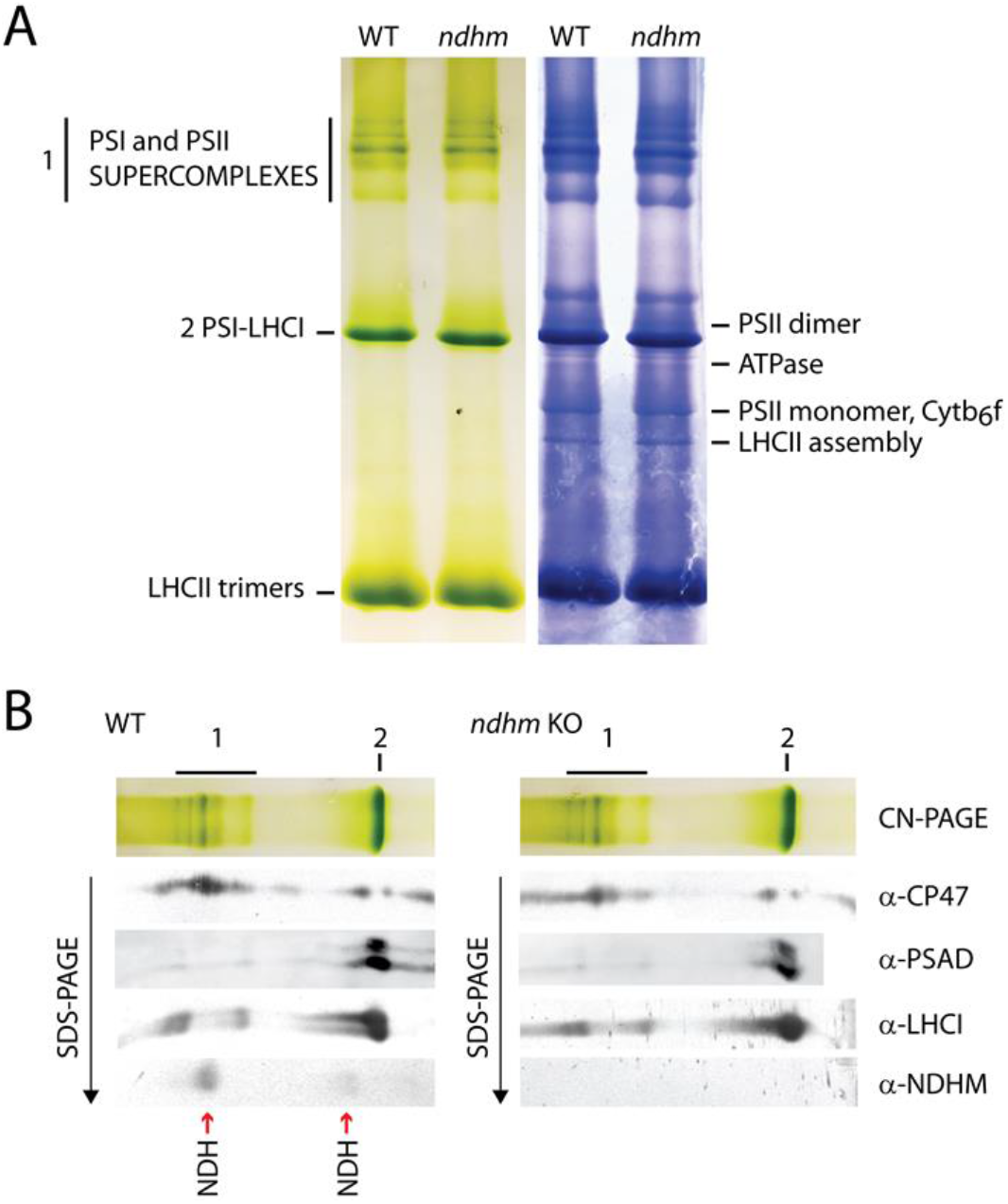
Native gel analysis of thylakoid membrane protein complexes. A) Protein complexes were solubilized from thylakoid membranes isolated from WT and *ndhm* KO protonema grown under control conditions. Protein complexes were separated by CN-PAGE and the known green bands were indicated on the left side while bands visible after Coomassie brilliant blue staining were indicated on the right side. B) The upper part of CN gel was further subjected to 2D SDS–PAGE and western blotting analysis using antibodies against CP47 for PSII, PSAD and LHCI for PSI and NDHM for NDH. Vertical red arrows indicate the positions of NDH as detected by the NDHM antibody. 1. PSI and PSII supercomplexes; 2. PSI.

### Response of the P. patens ndhm knock-out mutant to light

In order to assess the functional role of chloroplast NDH, a physiological characterization was performed analyzing the main photosynthetic parameters in plants exposed to increasing photosynthetic active radiation (Fig. S2). PSII quantum efficiency [Y(II)] upon exposure to increasing light intensities was indistinguishable in WT plants and in two independent clones of *ndhm* KO (Fig. S2). The Y(II), the NPQ level and the redox state of PQ (1-qL) did not show differences in the two genotypes independently from the light intensity (Fig. S3).

The efficiency of PSI [Y(I)] was instead slightly decreased in *ndhm* KO upon exposure to light (Fig. S2). This difference was attributable to an increased acceptor-side limitation [Y(NA)] in *ndhm* KO mosses as compared to WT (Fig. S2). It is interesting to notice that the impaired Y(I) translated into a lower electron transport rate of PSI (ETRI) over the one of PSII (ETRII) (Fig. S2). These results suggest a small but significant impairment of PSI activity in *ndhm* KO well consistent with results from analogous mutants in other plant species (Yamori *et al.*, 2015; Peltier *et al.*, 2016). In order to deeper analyze the higher Y(NA) of *ndhm* KO as compared to WT, we treated the plants for 8 minutes with two different intensities of constant actinic light (Fig. 3 and Fig. S3). Upon exposure to light comparable with the growth light intensity (50 μmol photons m^−2^ s^−1^), Y(I) was similar in both genotypes. The *ndhm* KO mutants showed a partially reduced donor-side limitation (Fig. 3B) and an increased acceptor-side limitation (Fig. 3C) compared to WT plants only immediately after the light was switched on. Acceptor side recovered to WT level but then increased again after few minutes of illumination. Similar results were obtained by treating the plants with stronger illumination (540 μmol photons m^−2^ s^−1^; Fig. 3D-F and Fig. S3) showing that *ndhm* KO mutation led to a higher Y(NA) limitation. Interestingly, Y(NA) was higher in *ndhm* KO than WT also when light was switched off and remained higher during the 10 minutes of dark recovery, both after 50 and 540 μmol photons m^−2^ s^−1^ of actinic light treatment (Fig. 3C and 3F). This suggested that in the absence of NDH activity, stromal acceptors are not readily oxidized as in WT after the light is switched off. As consequence Y(I) of *ndhm* KO is lower in the dark respect to WT plants (Fig. 3A and 3D). The failure to recover after several minutes in the dark suggests that this difference is likely attributable to a PSI photoinhibition suggesting that *ndhm* KO plants are more sensitive.

**Fig. 3.**
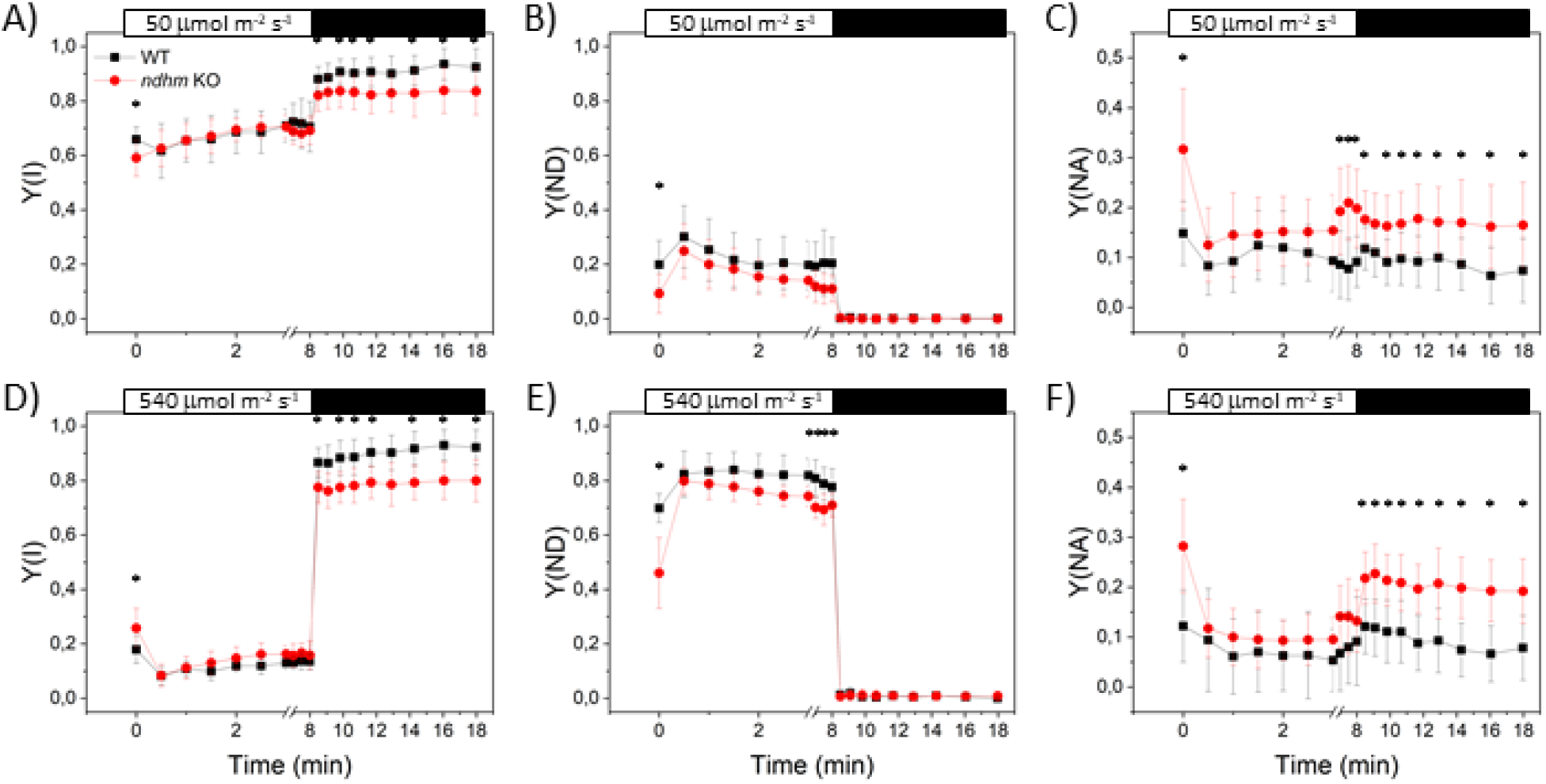
Effect of the NDHM depletion on P700 redox state. Y(I) (A, D), Y(ND) (B, E) and Y(NA) (C, F). White bars on top of each graph represent the interval in which actinic light was switched on and black bars represent the period of dark recovery. The actinic light intensity was 50 μmol photons m^−2^ s^−1^ for A-C and 540 μmol photons m^−2^ s^−1^ for D-F. For *ndhm* KO mutants, experiments were always performed using two independent lines and for clarity reasons the average value obtained from the two lines was presented. WT are represented by black squares and *ndhm* KO by red circles. Data represent average values ± sd, n = 9 - 14. (*P < 0.01; examined by One-way Anova).

### Interaction of NDH and pseudo-cyclic electron transport

*P. patens* was shown to have an intense PCEF dependent from the activity of FLV that transport electrons downstream PSI in the first seconds after an increase in light intensity, and their depletion results in PSI acceptor side limitation (Gerotto *et al.*, 2016). Since similarly also NDH is important to sustain PSI activity, its putative functional interaction with FLV-dependent pathways (Fig. 4A) was tested by introducing the *flva* KO mutation (Gerotto *et al.*, 2016) in *ndhm* KO plants (Fig. S4). The homologous recombination event in *flva ndhm* KO mutant plants was verified at the level of gDNA, while RT-PCR confirmed absence of *FLVA* expression (Fig. S4). Immunoblot analysis confirmed the impaired accumulation of NDHM and FLVA in all corresponding KO lines (Fig. 4). FLVB is also reduced by at least 90 % in double mutants as compared to WT as consequence of FLVA depletion. The *flva ndhm* KO mutants showed no change in PsaD, CP47, Cyt f and γ-ATPase as compared to reference lines (Fig. 4). As previously observed, the Y(I) of *flva* KO mutants was low as compared to WT when actinic light was switched on (Fig. 5A and 5D) and this was due to a high acceptor side limitation, (Y(NA), Fig. 5C and 5F). No significant difference was observed between *flva* KO and *flva ndhm* KO was detected suggesting that lack of FLV had a much larger impact on PSI activity effect than the lack of NDH regardless of the actinic light used (50 and 540 μmol photons m^−2^ s^−1^) (Fig 5 and Fig. S5). The estimation of total electron flow (TEF) using ECS signal in WT, *ndhm* KO, *flva* KO and *flva ndhm* KO showed that NDHM depletion had no detectable effect both in WT and *flva* KO background (Fig. S6). When plants where treated by DCMU to block PSII activity and measure solely the electron flow through PSI, *flva* KO and *flva ndhm* KO showed an increased transport as compared to WT and *ndhm* KO respectively (Fig. S6). Mutants carrying *ndhm* KO mutation did not show a significant reduction in CEF as compared to the corresponding backgrounds, either WT or *flva* KO (Fig. S5). Overall, these results clearly show that FLV activity has a major impact on electron transport and generation of pmf after light is switched on while the influence of NDH complex on light driven electron transport is instead quantitatively far less relevant.

**Fig. 4.**
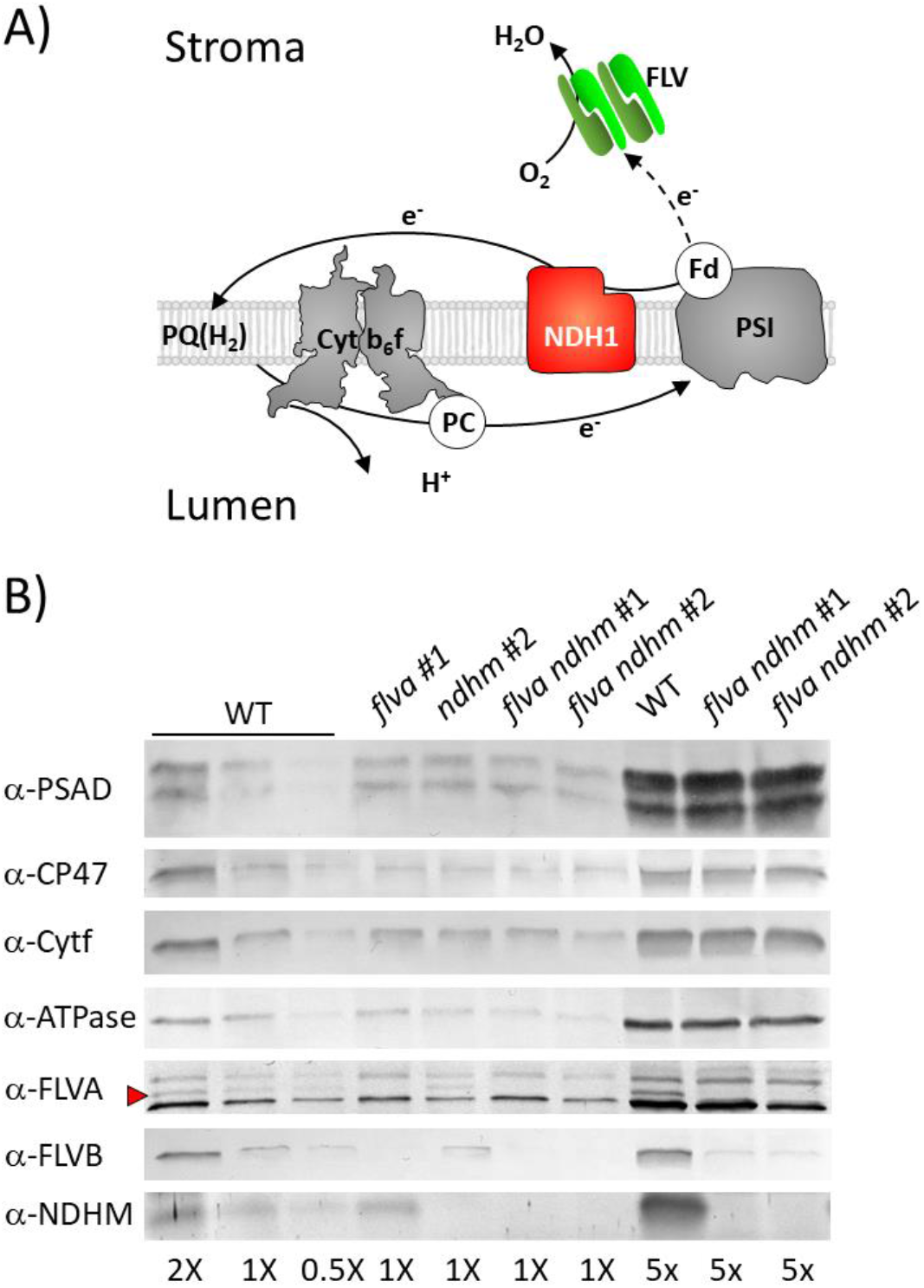
Isolation of *flva ndhm* double KO mutants. A) Schematic representation of NDH- and FLV-dependent electron transport pathways. B) Immunoblot analysis for detection of PSAD, CP47, Cytf, γ-ATPase, NDHM, FLVA and FLVB in WT, *flva* and *ndhm* single KO and *flva ndhm* double KO #1 and #2. 1X is equivalent to 1.5 μg of chlorophylls. 5X, 2X and 0.5X is respectively five, two and half times 1X.

**Fig. 5.**
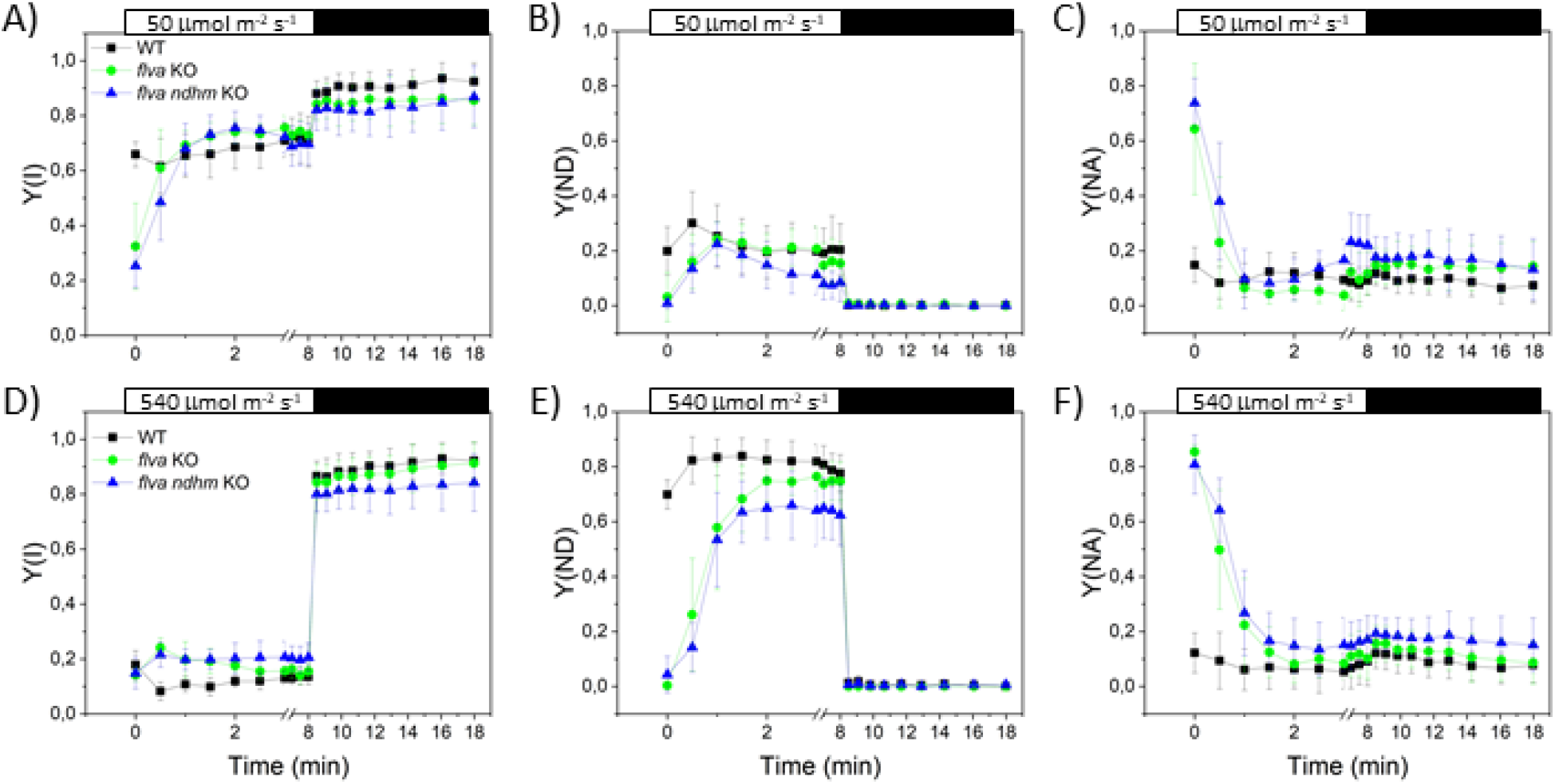
Effect of the FLVA depletion in a *ndhm* KO background on P700 redox state. Y(I) (A, D), Y(ND) (B, E) and Y(NA) (C, F). White bars on top of each graph represent the interval in which actinic light was switched on and black bars represent the period of dark recovery. The actinic light intensity was 50 μmol photons m^−2^ s^−1^ for A-C and 540 μmol photons m^−2^ s^−1^ for D-F. Data represent average values ± sd, n = 4 - 6. For *flva ndhm* KO mutants, experiments were always performed using two independent lines and for clarity reasons the average value obtained from the two lines was presented. All genotypes showed similar Fv/Fm, 0.82 ± 0.02 for WT, 0.83 ± 0.02 for flva KO and 0.83 ± 0.02 for flva ndhm KO. WT (black), *flva* KO (green) and flva *ndhm* KO (blue).

### NDH mutant lines are deficient in short-term response to changing light intensity

Previous data suggested an NDH complex activity during the dark-to-light transitions (Yamori *et al.*, 2016), even if the impact was less evident than for FLV. WT, single *ndhm* and *flva* KO and the double *ndhm flva* KO mutants were thus challenged by a reiterated exposure to saturating/limiting light cycles where 3 minutes of illumination with actinic light of 525 μmol photons m^−2^ s^−1^ were followed by 9 minutes of 25 μmol photons m^−2^ s^−1^ (Fig. 6). FLV proteins are a major sink of electrons during the low-to-high light transition (Gerotto *et al.*, 2016). So, in order to avoid that FLV electron transport would hide NDH complex activity, plants had been treated with longer and less frequent periods of high light choosing among the fluctuating light cycles previously tested for *P. patens* (Storti *et al.*, 2019). As expected, at each cycle the Y(I) was low (≈0.2) upon exposure to saturating-light period and it was high (≈0.8) when irradiance was limiting (Fig. 6A). In WT plants, this phenomenon depends on the limited availability of electron donors to PSI when light is in excess (Fig. 6C). The *ndhm* KO plants initially showed a similar phenotype as compared to WT, but with the reiteration of light cycles, an increase of acceptor side limitation was observed, causing a progressive reduction in Y(I) (Fig. 6A and E). Y(I) reduction also did not recover during the final dark phase, suggesting the onset of PSI photoinhibition rather than altered redox state in transporters. In the *flva ndhm* double KO, the responses of Y(I) and Y(NA) were more severe than those in the single mutants (Fig. 6,A and E), resulting also in a decline in Y(II) (Fig. 6B). Small but significant defects of NPQ activation were also accumulated by *flva ndhm* as compared to other genotypes (Fig. 6D) while their 1-qL was like those of *flva* KO (Fig. 6F). These results indicate that the re-iteration of light fluctuations has a cumulative effect and FLV and NDH complex cooperate for the protection of both photosystems in *P. patens* under fluctuating light.

**Fig. 6.**
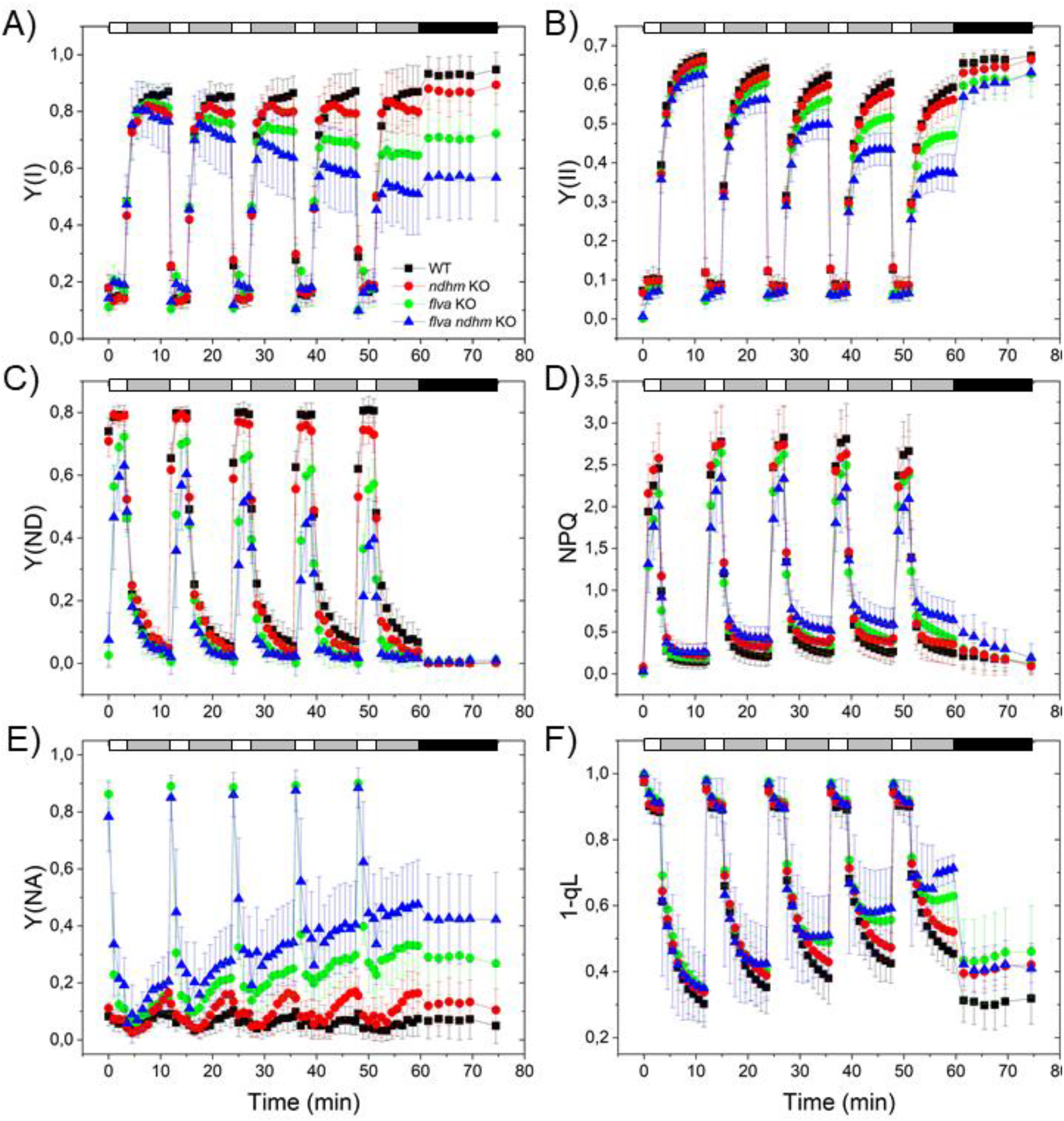
FLVA- and NDH-dependent electron transport protect photosystems under fluctuating light. Effect of fluctuating light on PSI and PSII: Y(I) (A), Y(II) (B), Y(ND) (C), NPQ (D), Y(NA) (E) and 1-qL (F) in WT (black squares), *ndhm* KO (red circles), *flva* KO (green circles) and *ndhm flva* KO (blue triangle), plants. Biological replicates, n = 8 ± sd for WT, 9 ± sd for *ndhm* KO, 7 ± sd for *flva* KO and 16 ± sd for *flva-ndhm* KO. At time 0, after 40 minutes of dark adaptation, plants were treated with saturating actinic light (535 μmol photons m^−2^ s^−1^) for 3 minutes followed by limiting actinic light (25 μmol photons m^−2^ s^−1^) 9 minutes. This 3+9 minutes cycle was repeated 4 more times. Differences between WT and mutant plants in the saturating/limiting light cycles were examined by One-way Anova, p values are reported in Table S3.

In order to evaluate whether the differences in photosynthetic performances had an impact on plant growth, we exposed WT, *flva* KO, *ndhm* KO, and double *flva ndhm* KO plants to different illumination regimes. Plants were first grown for 10 days in control conditions (CL, 50 μmol photons m^−2^ s^−1^) and then either kept in CL for the entire experiment or moved to other light conditions. There was no difference among genotypes under CL (Figure 7A) or under high light (HL, 500 μmol photons m^−2^ s^−1^) (Figure 7B). We also tested two different fluctuating light regimes (FL1 and FL2), delivering the same average light to plants (FL1 was ≈25 μmol photons m^−2^ s^−1^ for 9 min + ≈525 μmol photons m^−2^ s^−1^ for 3 min; FL2 was ≈25 μmol photons m-2 s-1 for 5 min and ~800 μmol photons m-2 s-1 for 1 min). Both fluctuating light regimes had a negative impact on plant growth, even in the case of WT plants. FL1 and FL2 did not affect *ndhm* KO plants as compared to WT while *flva* KO growth was reduced (Figure 7C and D). While under FL1 the impact of FLV was predominant, under FL2 instead the *flva ndhm* KO were more affected than single *flva* KO (Fig. 7D). This result suggests that in some conditions NDH complex and FLV-mediated electron transport have additional effect.

**Fig. 7.**
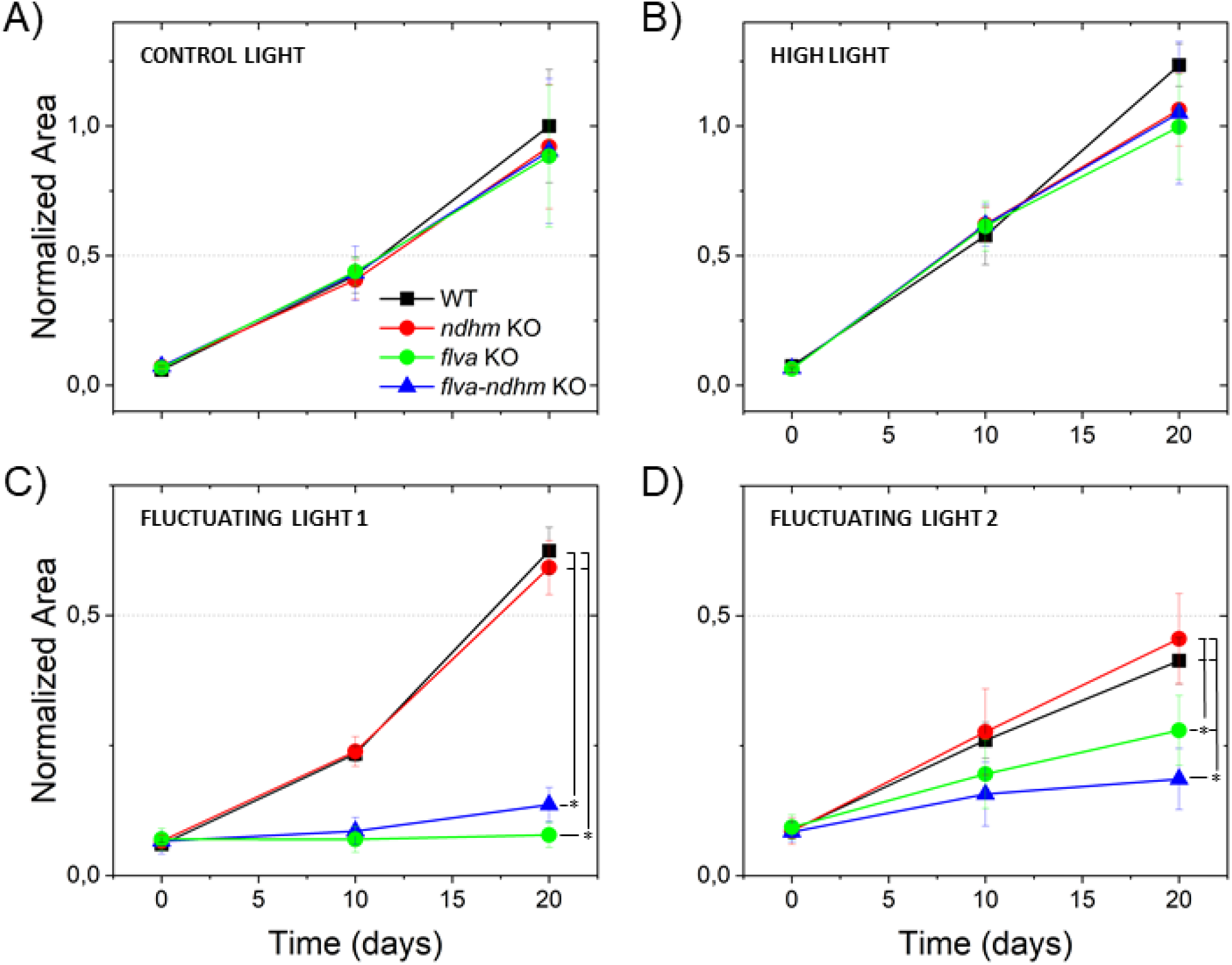
Phenotypes of wild-type (WT) and mutant plants grown under constant light and fluctuating light conditions. A) Under control light (50 μmol photons m^−2^ s^−1^) and B) under high light (500 μmol photons m^−2^ s^−1^) there were no differences in growth among genotypes. C) When light was switching between ≈25 μmol photons m^−2^ s^−1^ for 5 min and ≈800 μmol photons m^−2^ s^−1^ for 1 min, the *flva* KO and *flva ndhm* KO plants were smaller than WT and *ndhm* KO. D) When light was switching between ≈25 μmol photons m^−2^ s^−1^ for 9 min and ≈525 μmol photons m^−2^ s^−1^ for 3 min, the *flva ndhm* KO were smaller than *flva* KO that were smaller than *ndhm* KO and WT plants. For each sample, the area was normalized to the area of 20-day-old WT plants grown in control light conditions. The difference in relative growth rate is significant between WT/*ndhm* KO and *flva* KO/*ndhm* KO (C and D) and between *flva* KO and *flva ndhm* KO (D) (*p < 0.01; statistical analyses by Anova). From 3 to 14 biological replicates are here reported.

## DISCUSSION

It was recently proposed that NDH can form a supercomplex with one single PSI in *P. patens* (Kato *et al.*, 2018), instead of binding two PSI trough the antenna linkers LHCA5 and LHCA6 as in Angiosperms (Peng *et al.*, 2009; Kouřil *et al.*, 2014). Indeed, since *P. patens* genome contains LHCA5 but not a LHCA6 homologue (Alboresi *et al.*, 2008), only one PSI antenna should be available to bind NDH. Results presented here on *ndhm* KO of *P. patens* are consistent with this hypothesis. In fact, in WT plants NDHM was detected in a region of the gel compatible with the apparent molecular weight of a PSI-NDH supercomplex. However, we could not demonstrate the comigration of NDH with PSI, leaving open alternative possibilities, such as either this PSI-NDH supercomplex is unstable in the conditions used for solubilization and CN-PAGE or NDH is involved in the formation of a dimer or of another heterocomplex (Fig. 2B). However, since most NDHM is detected in this putative supercomplex, 2D-immunoblot results suggest that if a PSI-NDH complex is indeed present it represents a very small fraction of the whole PSI population that would be fully consistent with the observation of a reduced impact of the mutation on the total PSI electron transport capacity.

*P. patens* plants defective in NDH complex did not show any major growth or developmental phenotype if compared to WT (Fig. 1), consistently to what was observed in several other organisms like *Marchantia, Nicotiana, Arabidopsis* and *Oryza* (Endo *et al.*, 1999; Ishikawa *et al.*, 2008; Yamori *et al.*, 2015). On the contrary NDH-mediated CEF might be essential in C4 plants, as demonstrated by NDHN suppression in *Flaveria bidentis* (Ishikawa *et al.*, 2016) and might have a key roles in the development of specific plant organs, as suggested by transposon-tagged tomato plants in which the loss of NDHM expression impairs tomato fruit ripening (Nashilevitz *et al.*, 2010). This functional heterogeneity, together with the variable distribution of NDH complex among photosynthetic organisms (Ruhlman *et al.*, 2015), suggest that NDH complex physiological role during plastid evolution specialized over time (de Vries *et al.*, 2016) but in some cases it has been juxtaposed or replaced by other mechanisms.

The analysis of photosynthetic parameters of *ndhm* KO plants at different light intensities revealed that NDH complex mutation has a small impact on Y(NA) during the dark-to-light transitions (Fig. S2 and Fig. 3), both under control light (50 μmol photons m^−2^ s^−1^) and under high light (540 μmol photons m^−2^ s^−1^) conditions. The phenotype observed at the level of PSI acceptor side is congruent with previous works demonstrating that CEF is important during the first seconds of photosynthetic activity in C3 plants (Joliot and Joliot, 2006).

When the NDH inactivation was combined with depletion of FLV it emerged that the latter have a larger role in electron transport capacity than NDH complex. FLV are responsible for the transport of a high number of electrons in the first seconds after light is switched on and their role cannot be compensated by the low electron transport capacity of NDH complex (Shikanai, 2016*a*; Strand *et al.*, 2017) (Fig. 6, Fig. 7 and Fig. S5). This observation is supported by the biochemical evidences presented here, with likely only a small fraction of PSI involved in super-complexes with NDH. If this is the case, in WT plants, when PSI is fully involved in transporting electrons, NDH activity is not detectable behind the one of FLV proteins.

The small impact of NDH on photosynthetic activity is however amplified when fluctuating light cycles are reiterated and the difference between WT and *ndhm* KO increases progressively (Fig. 6). At each repetition the difference in Y(NA) limitation between *ndhm* KO and WT plants increased (Fig. 6E), ultimately leading to a significant impairment in PSI yield (Fig. 6A). Interestingly, although both FLV and NDH influence Y(NA), the effect of FLV is more evident in the low-to-high light intensity switch, while the effect of NDH complex increases during the low light period. The repetition of the cycles also indirectly affects PSII activity, leading to a more pronounced reduction of PQ pool in *ndhm* KO plants than in WT (Fig. 6). The role of NDH complex is well compensated by other electron transport pathways when plants are grown in constant conditions of a growth chamber. Cyclic light experiment suggests that NDH complex physiological role could be much more relevant in natural environments, where the fluctuations in light intensities are frequent. In this case, NDH-dependent CEF pathway could provide higher flexibility to the chloroplast electron transport chain. Interestingly, a functional redundancy and crosstalk between FLV and NDH-1 complex were recently observed also in *Synechocystis sp.* PCC 6803 for the dynamic coordination of PSI oxidation and for photoprotection under variable CO_2_ and light availability (Nikkanen *et al.*, 2020).

The physiological role in fluctuating light is apparently in contrast with the observations of a limited influence on photosynthetic electron transport. To reconcile this apparent discrepancy it should be underlined that while NDH is not abundant and active enough to have a major impact on electron transport when this is under full activity, the picture could instead be different in the dark or under limiting light intensities, when light-driven electron transport is reduced and where NDH activity could become physiologically more impactful. Indeed, we observed that in *ndhm* KO PSI acceptor side limitation increased also during limiting light condition and in the dark at the end of the kinetic, suggesting that NDH could be active in maintaining PSI acceptor side oxidized after the light is switched off. The effect is visible when light is increased again, thus, it would not be due to the NDH electron transport ability but rather to the fact that PSI in the absence of NDH is already partially reduced, limiting its acceptor capacity and leading to PSI photoinhibition. A functional role of NDH complex in the dark period of fluctuating light treatments was already reported in the case of *O. sativa* (Yamori *et al.*, 2016) and *A. thaliana* (Strand *et al.*, 2017). In both cases, WT plants decreased the electron flow to PSI during the light phase, which seemed to be essential to prevent the over-reduction of PSI and to keep the reduction level of the entire electron transport chain low enough during the subsequent dark or low-light phase (Yamori *et al.*, 2016; Strand *et al.*, 2017).

It should be asked why this could be advantageous for plants. PSI is known to be stable under saturating light conditions, thanks to P700^+^ that is a good quencher of excess energy. On the contrary PSI is sensitive to an excess of electrons that, once accumulated at the acceptor side, can generate reactive oxygen species because of the presence of Fe-S clusters (Sonoike *et al.*, 1995; Tiwari *et al.*, 2016). PSI under strong illumination is normally found limited at the donor side avoiding ROS production and PSI photoinhibition (Larosa *et al.*, 2018). After the light is switched off, NDH complex contributes to keeping PSI not limited at the acceptor side and reducing the probability of an acceptor side limitation when light is back on. There are also biochemical implications, since NDH participates to the regulation of the electron flow that leads to the formation of NADPH pools in the light. Then NADPH is directly used as cofactor in enzymatic reactions and it is also produced during darkness from sugars by the oxidative pentose phosphate pathway (Cejudo *et al.*, 2019). The *ndhm* KO lines redox imbalance in low light and in the dark could be of physiological importance to redox-dependent reactions in the chloroplast and the light reactions of photosynthesis.

## SUPPLEMENTARY DATA

Table S1. Primers employed for *ndhm* KO generation and screening.

Table S2. Pigment composition of *Physcomitrella patens* WT and *ndhm* KO plants.

Table S3. Statistical analysis by one-way Anova of the experiment reported in Figure 6

Fig. S1. Isolation of *ndhm* KO mutants of *Physcomitrella patens*.

Fig. S2. Effect of NDHM deletion on the photosynthetic activity of *Physcomitrella patens*.

Fig. S3. Effect of the NDHM depletion on photosystem II activity.

Fig. S4. Isolation of double *flva ndhm* KO mutants from the single *ndhm* KO #2 background.

Fig. S5. Effect of the FLVA depletion in a *ndhm* KO background on photosystem II activity.

Fig. S6. Photosynthetic electron transport in *P. patens* plants.

## ACKNOWLEDGMENTS

TM and AA planned and designed the research. MS performed most of the experiments and analyzed the data. MPP isolated the mutants. MS, MPP, performed experiments. MS, AA analyzed the data. TM, MS and AA wrote the manuscript, which all authors revised and approved. Anti-NDHH antibody was kindly provided by Dominique Rumeau (CEA Cadarache, France). Caterina Gerotto and Marco Armellin for help in preliminary experiments. This study was supported by funding from the Università degli Studi di Padova, Dipartimento di Biologia [BIRD173749/17].

## ABBREVIATIONS

gDNA: genomic DNA
PSII: photosystem II
Cyt b_6_f: cytochrome b_6_f
PSI: photosystem I
PQH_2_: plastoquinone
LEF: linear electron flux
CBB: Calvin Benson Bassham
CEF: cyclic electron flux
LHC: light harvesting complex
ETR I: electron transport rate through photosystem I
ETR II: electron transport rate through photosystem II

