## Supplementary Tables and Figures for "The activity of chloroplast NADH dehydrogenase-like complex influences the photosynthetic activity of the moss *Physcomitrella patens*"

**Table S1. Primers employed for *ndhm* KO and *flva* KO generation and screening.** The table reports the primers used to generate and characterize *Physcomitrella patens ndhm* KO lines and *ndhm flva* double KO lines. Primers were designed to disrupt *NDHM* coding sequence with the insertion of the bleomycin resistance cassette (ble). Genomic DNA from WT strain was used as a template to amplify selected homologous regions (P1 + P2 and P3 + P4). All the PCR products were cloned into BZRf vector for moss transformation using the restriction enzymes (REs) indicated. Moss protoplasts were transformed with *NDHM* KO construct linearized with PmlI and PacI. After transformation and two rounds of selection on zeocin, resistant lines were evaluated for correct homologous recombination using P5 + P6 (Left Border, LB) and P7 + P8 (Right Border, RB) primers. In order to disrupt *FLVA* gene in *ndhm* KO plants, we used the same vector and protocol that previously allowed the isolation of *flva* KO (Gerotto *et al.*, 2016) and *flva pgrl1* KO (Storti *et al.*, 2019). After transformation and two rounds of selection on hygromycin-B, resistant lines were screened using FLVA_5 + P6 (Left Border, LB) and FLVA_6 + 35S terminator (Right Border, RB) primers. Primers NDHM_F, NDHM_R, FLVA_7 and FLVA_8 were used on cDNA libraries to verify *NDHM* expression by RT-PCR. ACTIN2_F and ACTIN2_R were used to control the quality of genomic DNA and cDNA templates. (*) indicates primers already used for the isolation of *flva* KO and *flva pgrl1* KO (Gerotto *et al.*, 2016; Storti *et al.*, 2019).

| **Primer name** | **Gene** | **Sequence** | **Use (RE)** |
|---|---|---|---|
| P1 | *NDHM* | atCACGTGCAGCTGCAACAAGTACCCAG | Vector design (PmlI) |
| P2 | *NDHM* | atCTCGAGAATTCGCACATGACGAGTCG | Vector design (XhoI) |
| P3 | *NDHM* | atGTTAACTTGGACTGTAGGGTGCTGAAC | Vector design (HpaI) |
| P4 | *NDHM* | atTTAATTAATTACAAGCCAGCAAAGCAAA | Vector design (PacI) |
| P5 | *NDHM* | TTGGAAGTCTGTTCACGCTTT | KO screening |
| P6 | *35S promoter* | GTGTCGTGCTCCACCATGT | KO screening |
| P7 | ble | CCCCGCTTAAAAATTGGTAT | KO screening |
| P8 | *NDHM* | TTCTGCCAATAGGATGTGAGG | KO screening |
| FLVA_5 (*) | *FLVA* | CACACTCTACATCGAAGCCTCA | KO screening |
| FLVA_6 (*) | *FLVA* | GCTAAGCGCAGCAACACTTT | KO screening |
| P9 | *35S terminator* | CGCTGAAATCACCAGTCTCTCT | KO screening |
| NDHM_F | *NDHM* | AGTGTCCTCCGCTTTTCTCA | RT-PCR |
| NDHM_R | *NDHM* | CTCCGTCAAATCTGCACCTG | RT-PCR |
| FLVA_7 (*) | *FLVA* | TTTGGCTCTTTCGGGTGGAG | RT-PCR |
| FLVA_8 (*) | *FLVA* | GACGGTTTTCGCCAGGTTTG | RT-PCR |
| ACTIN2_F | *ACTIN2_F* | GCGAAGAGCGAGTATGACGAG | RT-PCR |
| ACTIN2_R | *ACTIN2_R* | AGCCACGAATCTAACTTGTGATG | RT-PCR |

**Table S2. Pigment composition of *Physcomitrella patens* WT and *ndhm* KO plants.** Chlorophyll a/b and chlorophyll/carotenoid ratios (mol/mol) of protonema with SDs are shown (n = 5 to 8 for each genotype).

| | **Chl a/b** | **chl/car** |
|---|---|---|
| WT | 2.56 ± 0,11 | 3,85 ± 0,43 |
| *ndhm#1* KO | 2,62 ± 0,15 | 4,08 ± 0,47 |
| *ndhm#2* KO | 2,63 ± 0,13 | 3,67 ± 0,33 |

**Table S3. One-way Anova of the photosynthetic characterization of WT and mutant lines reported in Figure 6.** All points of each cycle were considered for statistical analysis, all comparisons with p < 0,01 are highlighted in green.

| | | Y(I) | Y(NA) | Y(ND) | Y(II) | NPQ | qL |
|---|---|---|---|---|---|---|---|
| Cycle 1 | WT vs *ndhm* KO | 1,81E-02 | 2,96E-02 | 1,81E-01 | 2,49E-01 | 9,86E-03 | 2,18E-01 |
| | WT vs *flva* KO | 2,70E-01 | 1,93E-06 | 6,45E-01 | 1,82E-01 | 7,58E-01 | 3,18E-02 |
| | WT vs *flva ndhm* KO | 2,85E-02 | 4,11E-09 | 5,10E-01 | 4,70E-03 | 3,08E-02 | 4,48E-01 |
| | *ndhm* KO vs *flva* KO | 2,64E-01 | 3,23E-02 | 8,99E-02 | 8,20E-01 | 1,60E-02 | 2,53E-01 |
| | *ndhm* KO vs *flva ndhm* KO | 9,82E-01 | 1,67E-05 | 4,65E-02 | 1,02E-01 | 4,18E-01 | 8,02E-01 |
| | *flva* KO vs *flva ndhm* KO | 3,06E-01 | 1,27E-02 | 8,77E-01 | 1,86E-01 | 4,42E-02 | 2,52E-01 |
| Cycle 2 | WT vs *ndhm* KO | 9,02E-02 | 2,79E-05 | 9,14E-01 | 1,88E-01 | 9,01E-03 | 1,31E-01 |
| | WT vs *flva* KO | 9,61E-03 | 7,58E-20 | 5,38E-02 | 9,25E-02 | 7,14E-02 | 1,15E-01 |
| | WT vs *flva ndhm* KO | 1,01E-04 | 1,06E-21 | 5,08E-03 | 7,57E-05 | 1,04E-04 | 4,28E-01 |
| | *ndhm* KO vs *flva* KO | 3,36E-01 | 4,34E-08 | 7,63E-02 | 6,69E-01 | 3,04E-01 | 8,14E-01 |
| | *ndhm* KO vs *flva ndhm* KO | 1,41E-02 | 3,46E-14 | 8,40E-03 | 8,14E-03 | 3,01E-01 | 6,07E-01 |
| | *flva* KO vs *flva ndhm* KO | 1,31E-01 | 5,75E-04 | 4,75E-01 | 3,52E-02 | 2,27E-02 | 5,12E-01 |
| Cycle 3 | WT vs *ndhm* KO | 1,36E-01 | 8,04E-07 | 4,67E-01 | 1,50E-01 | 1,35E-02 | 5,71E-02 |
| | WT vs *flva* KO | 1,59E-04 | 3,03E-29 | 2,66E-03 | 1,11E-02 | 6,40E-02 | 2,19E-03 |
| | WT vs *flva ndhm* KO | 2,49E-09 | 2,66E-31 | 2,49E-05 | 3,49E-09 | 4,01E-06 | 1,64E-02 |
| | *ndhm* KO vs *flva* KO | 1,28E-02 | 1,51E-14 | 2,43E-02 | 2,46E-01 | 4,07E-01 | 1,49E-01 |
| | *ndhm* KO vs *flva ndhm* KO | 5,27E-07 | 4,57E-23 | 7,11E-04 | 5,07E-06 | 6,36E-02 | 3,73E-01 |
| | *flva* KO vs *flva ndhm* KO | 5,38E-03 | 7,60E-06 | 3,55E-01 | 8,00E-04 | 2,73E-03 | 7,31E-01 |
| Cycle 4 | WT vs *ndhm* KO | 1,91E-01 | 2,17E-08 | 1,10E-01 | 1,53E-01 | 5,35E-02 | 1,29E-02 |
| | WT vs *flva* KO | 1,24E-07 | 9,63E-37 | 2,18E-05 | 2,20E-04 | 4,03E-02 | 4,23E-06 |
| | WT vs *flva ndhm* KO | 1,44E-16 | 3,75E-40 | 1,85E-08 | 1,20E-16 | 9,03E-07 | 9,74E-06 |
| | *ndhm* KO vs *flva* KO | 1,07E-05 | 1,75E-20 | 9,65E-03 | 1,92E-02 | 9,66E-01 | 8,84E-03 |
| | *ndhm* KO vs *flva ndhm* KO | 7,59E-15 | 6,46E-31 | 1,56E-04 | 9,16E-13 | 4,96E-03 | 6,08E-03 |
| | *flva* KO vs *flva ndhm* KO | 3,30E-05 | 4,21E-07 | 3,18E-01 | 1,88E-07 | 1,57E-03 | 5,78E-01 |
| Cycle 5 | WT vs *ndhm* KO | 3,05E-01 | 7,53E-08 | 3,79E-02 | 1,28E-01 | 1,72E-01 | 4,02E-04 |
| | WT vs *flva* KO | 6,44E-12 | 6,57E-37 | 1,78E-06 | 8,68E-08 | 1,21E-02 | 5,96E-11 |
| | WT vs *flva ndhm* KO | 3,61E-25 | 1,37E-40 | 2,04E-10 | 6,80E-26 | 1,04E-08 | 1,42E-14 |
| | *ndhm* KO vs *flva* KO | 5,11E-11 | 1,27E-20 | 8,92E-03 | 7,16E-05 | 2,82E-01 | 3,58E-05 |
| | *ndhm* KO vs *flva ndhm* KO | 1,64E-25 | 3,78E-31 | 6,69E-05 | 8,96E-22 | 1,44E-05 | 3,67E-09 |
| | *flva* KO vs *flva ndhm* KO | 5,74E-08 | 4,21E-07 | 1,65E-01 | 4,34E-13 | 4,70E-04 | 1,07E-02 |

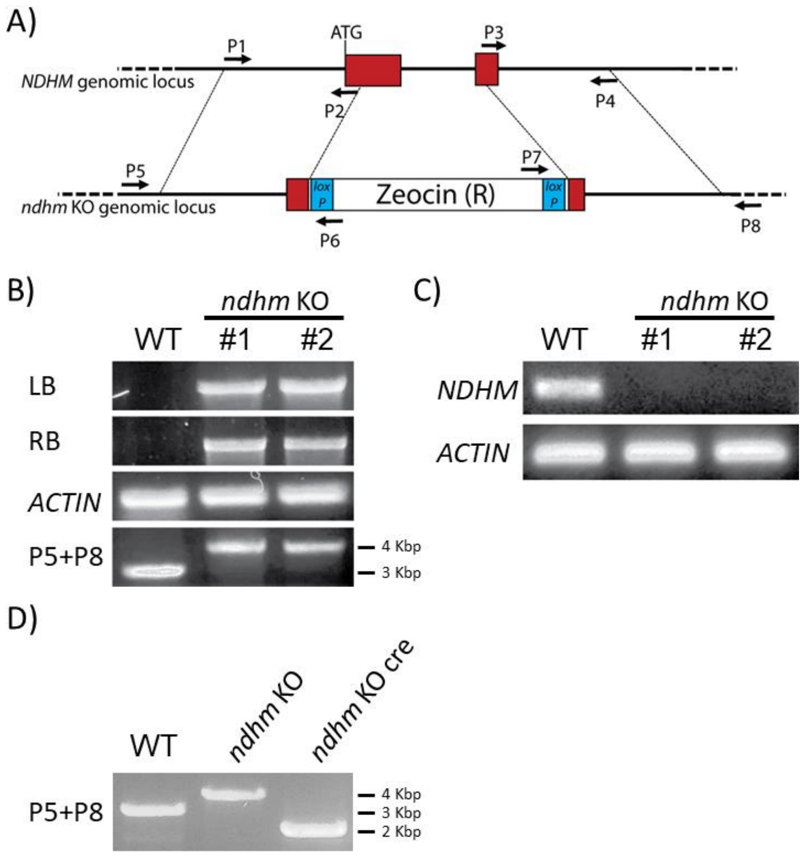


**Fig. S1. Isolation of *ndhm* KO mutants of *Physcomitrella patens*.** A) Schematic representation of the *NDHM* gene and corresponding *ndhm* KO. Red rectangles indicate exons and black arrows represent the primers used for cloning (from P1 to P4) and screening (from P5 to P8) procedures. After homologous recombination, the central part of the gene is substituted by the resistance cassette. Dotted lines indicate the regions of the gene used to perform homologous recombination. Blue boxes indicate the LoxP sites used for Cre/lox recombination and excision of the cassette conferring zeocine resistance. B) Confirmation of homologous recombination event in the *NDHM* locus. Left (LB) and right borders (RB) were respectively amplified by P5/P6 and P7/P8. Positive results indicated that the resistance cassette was inserted in the target locus. *ACTIN* primers were used as quality control for gDNA. Primers P5 and P8 amplified the WT locus (~3 Kbp) and *ndhm* KO locus with one single insertion (≈4 Kbp). Multiple insertions in the target locus could not be amplified by PCR, therefore the corresponding lines were discarded to focus on single insertion lines #1 and #2. C) RT-PCR analysis for the detection of *NDHM* and *ACTIN* transcripts in WT and two independent putative KO mosses. D) Analysis of the recombinant locus after transient expression of the Cre recombinase in *ndhm* KO plants. Primers P5 and P8 amplified WT, *ndhm* KO and *ndhm* KO cre locus lacking the resistance cassette (≈2 Kbp). Primer sequences is reported in Table S1.

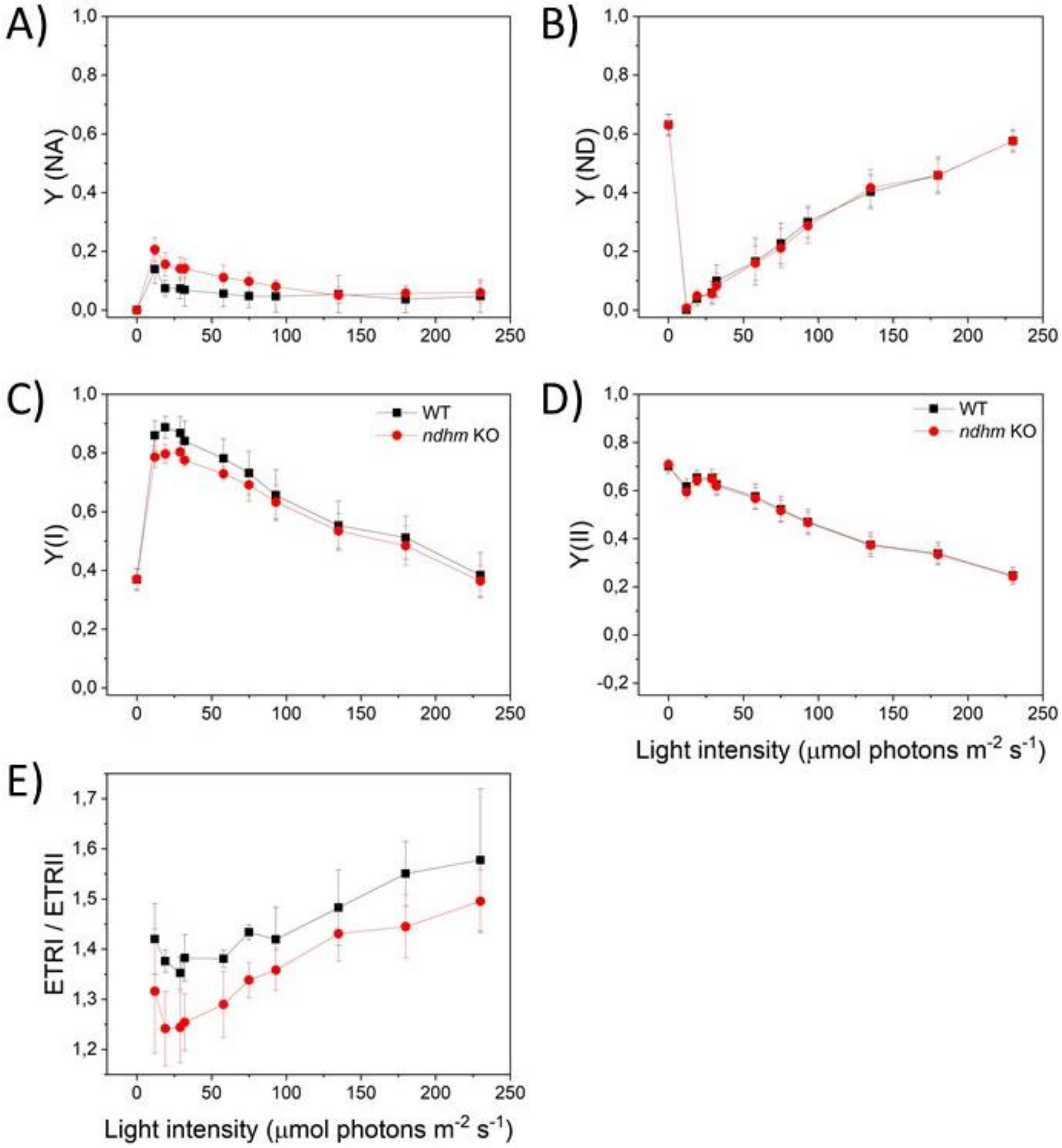


**Fig. S2. Effect of NDHM deletion on the photosynthetic activity of *Physcomitrella patens*.** Light-intensity dependence of (A) Y(NA), (B) Y(ND), (C) Y(I), (D) Y(II) and (E) ETRI/ETRII in WT (black squares) and two independent lines of *ndhm* KO (red circles), biological replicates n=3-6 ± sd. The maximum PSII efficiency was 0.83 ± 0.02 in WT and 0.84 ± 0.02 in *ndhm* KO. For *ndhm* KO mutants, experiments were always performed using two independent lines and for clarity reasons the average value obtained from the two lines was presented.

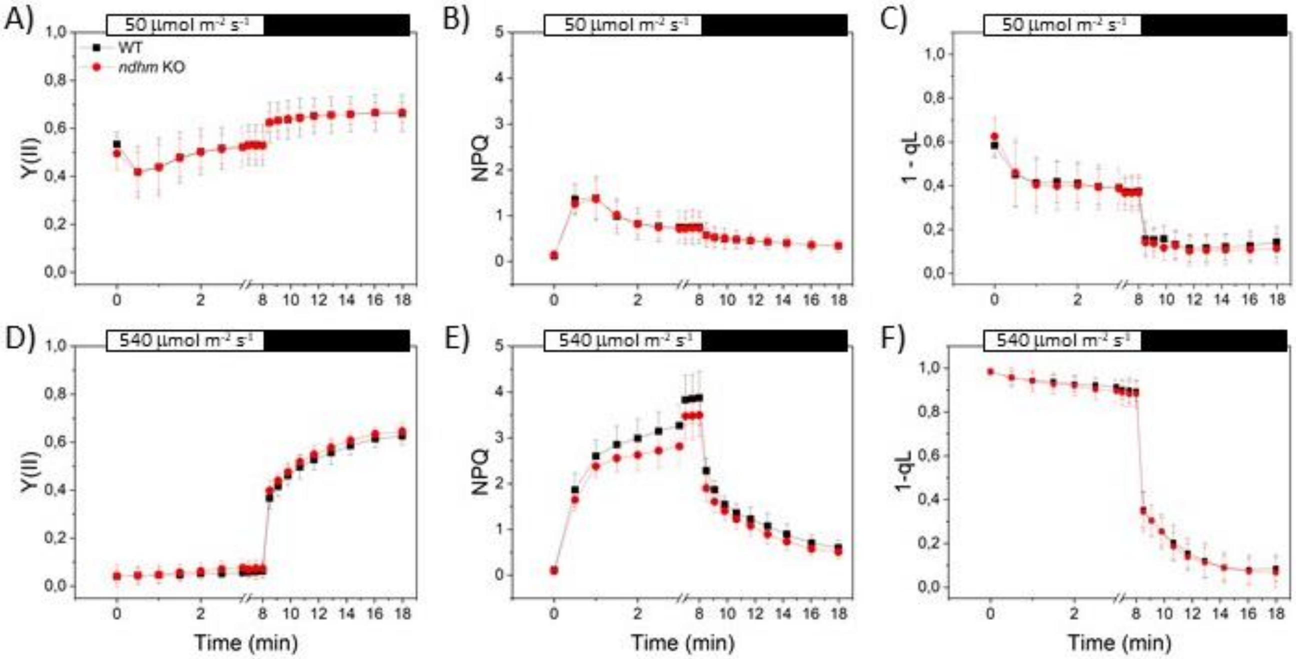


**Fig. S3. Effect of the NDHM depletion on photosystem II activity.** Chlorophyll fluorescence was measured to calculate time course of Y(II) (A, D), NPQ (B, E) and 1-qL (C, F). White and black bars on top of each graph represent the interval in which actinic light was switched respectively on and off. Actinic light intensity was 50 µmol photons $m^{-2}$ $s^{-1}$; (A-C) and 540 µmol photons $m^{-2}$ $s^{-1}$ (D-F). For *ndhm* KO mutants, experiments were always performed using two independent lines and for clarity reasons the average value obtained from the two lines was presented. WT (black) and *ndhm* KO (red). Data represent mean values ± sd, n = 4 - 10.

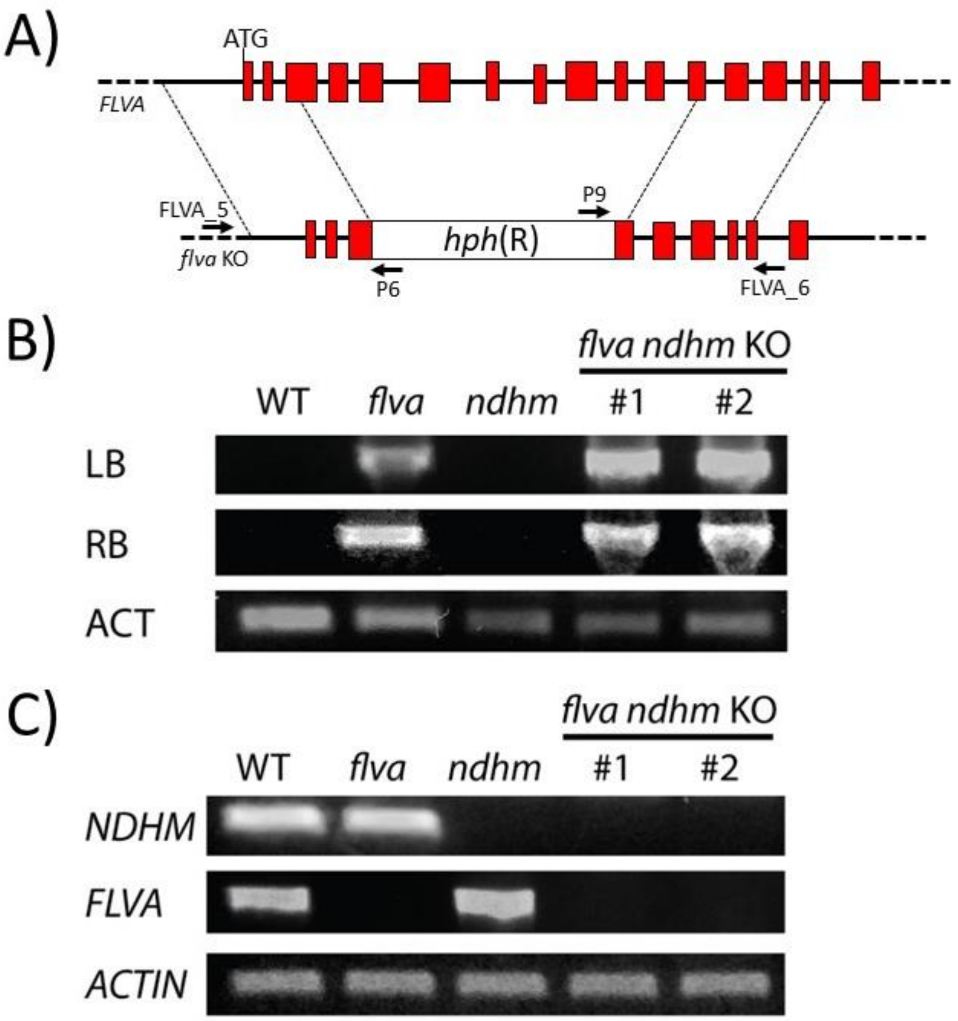


**Fig. S4. Isolation of double *flva ndhm* KO mutants from the single *ndhm* KO #2 background.** A) Schematic representation of the *FLVA* gene and corresponding *flva* KO locus. Red rectangles indicate exons and black arrows represent the primers (FLVA_5, P6, P9 and FLVA_6) used for screening procedures. After homologous recombination, the central part of the gene is substituted by the resistance cassette. Dotted lines indicate the regions of the gene used to perform homologous recombination. B) Confirmation of homologous recombination event in the *FLVA* locus of *ndhm* KO mutant. LB and RB stand respectively for left and right borders obtained as previously described for the isolation of *flva* KO single mutants (Gerotto *et al*., 2016). C) RT-PCR analysis for the detection of *NDHM*, *FLVA* and *ACTIN* transcripts in WT, *flva* KO, *ndhm* KO and two putative *flva ndhm* KO mosses. Primer sequences are reported in Table S1.

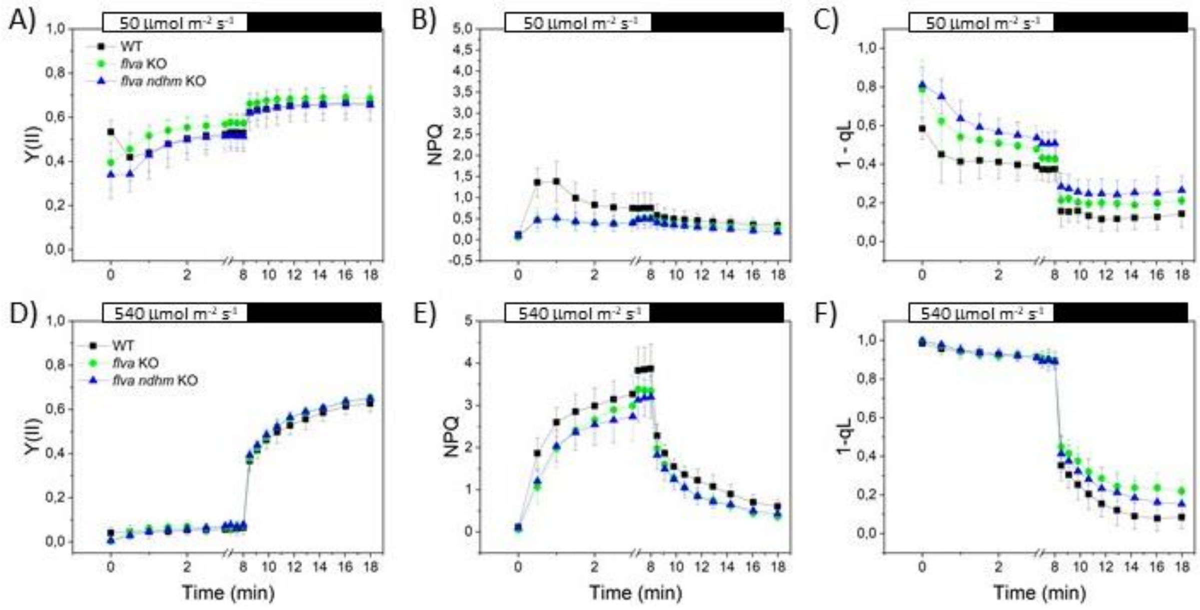


**Fig. S5. Effect of the FLVA depletion in a *ndhm* KO background on photosystem II activity.** Chlorophyll fluorescence was measured to calculate time course of Y(II) (A, D), NPQ (B, E) and 1-qL (C, F). White and black bars on top of each graph represent the interval in which actinic light was switched respectively on and off. Actinic light intensity was 50 µmol photons $m^{-2}$ $s^{-1}$; (A-C) and 540 µmol photons $m^{-2}$ $s^{-1}$ (D-F). For *flva ndhm* KO mutants, experiments were always performed using two independent lines and for clarity reasons the average value obtained from the two lines was presented. WT (black), *flva* KO (green) and *flva ndhm* KO (blue). Data represent mean values ± sd, n = 4 - 10.

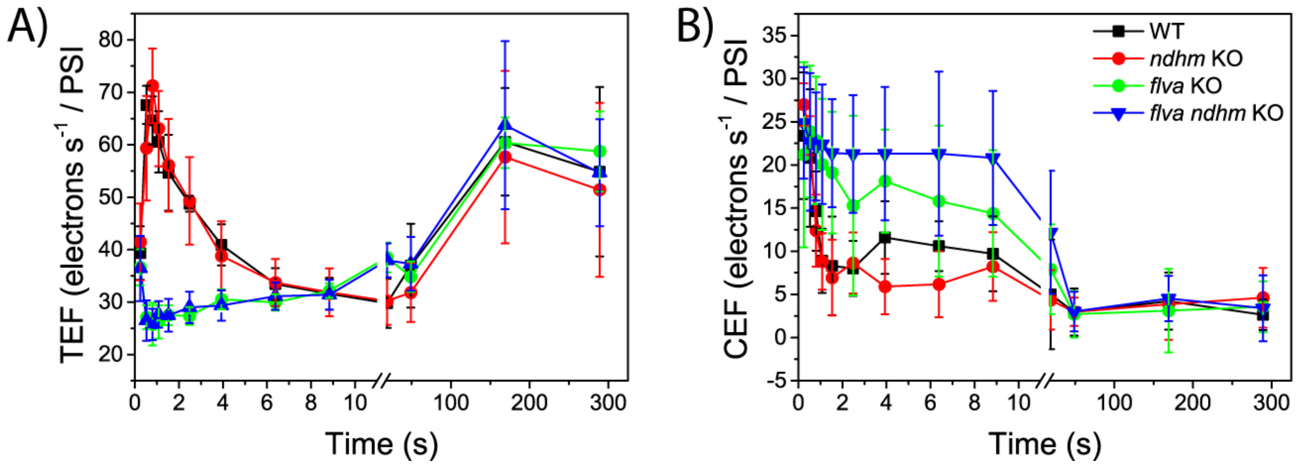


**Fig. S6. Photosynthetic electron transport in *P. patens* plants.** WT and mutant lines were grown in control light for 10 days. A) Total photosynthetic electron flow (TEF) and cyclic electron flow (CEF) measured *in vivo* in WT (black squares), *ndhm* KO (red circles), *flva* KO (green circles) and *flva ndhm* KO (blur triangles) at 940 μmol photons $m^{-2} \cdot s^{-1}$ actinic light, calculated from electrochromic shift signal. Electron transport rate values are normalized to xenon-induced PSI turnovers. Data presented are averages ± sd; n = 6-12.
